# Predictors of zoonotic potential in helminths

**DOI:** 10.1101/2021.03.28.437423

**Authors:** Ania A. Majewska, Tao Huang, Barbara Han, John M. Drake

## Abstract

Helminths are parasites that cause disease at considerable cost to public health and present a risk for emergence as novel human infections. Although recent research has elucidated characteristics conferring a propensity to emergence in other parasite groups (e.g. viruses), the understanding of factors associated with zoonotic potential in helminths remains poor. We applied an investigator-directed learning algorithm to a global dataset of mammal helminth traits to identify factors contributing to spillover of helminths from wild animal hosts into humans. We characterized parasite traits that distinguish between zoonotic and non-zoonotic species with greater than 88% accuracy. Results suggest that helminth traits relating to transmission (e.g. definitive and intermediate hosts) and geography (e.g. distribution) are more important to predicting zoonotic species than morphological or epidemiological traits. Whether or not a helminth causes infection in companion animals (cats and dogs) is the most important predictor of propensity to cause human infection. Finally, we identified helminth species with high modeled propensity to cause zoonosis (over 70%) that have not previously been deemed to be of risk. This work highlights the importance of prioritizing studies on the transmission of helminths that infect pets and points to the risks incurred by close associations with these animals.

## Introduction

Understanding the factors that contribute to the emergence of novel infectious diseases is a central concern to global public health [1]. Since most outbreaks of novel pathogens among humans are due to spillover from animal hosts [2-4], identifying factors associated with the propensity for transmission to humans is of high priority. Research in this area is particularly urgent because the rate of human-wildlife contacts is increasing with changes to natural landscapes and global climate [5], providing ample opportunities for human exposure to novel hosts and pathogens [6, 7]. Identifying species that are potentially parasitic or pathogenic in humans (i.e., those with high *zoonotic potential*) would enhance our understanding of the factors underpinning spillover transmission from animal reservoirs, and enable preemptive approaches to disease control.

One approach to evaluating zoonotic potential is to analyze pathogen and host traits [e.g. 8]. Particularly, features distinguishing zoonotic from non-zoonotic parasites and their reservoir host species can be used to predict which species are most likely to present high risk of zoonotic exposure to people [9]. For example, work by Han, Schmidt [10] identified ‘fast’ life history strategy (short-lived, short generation time) as a key predictor of the rodent species most likely to be reservoirs of novel zoonotic pathogens. Trait analysis of zoonotic viruses revealed that viruses which can replicate in cytoplasm are more likely to infect humans [11] and viruses which infect nonhuman primates predict the transmissibility of a virus between humans [12]. Patterns in genome sequences of viruses have also yielded predictions on which hosts are likely to be reservoirs of zoonosis and which arthropods are likely to be their vectors [13]. These findings are of scientific interest concerning the current theoretical debate about why some parasite species are more prone to spillover [14-16].

Parasitic helminths are a group of parasites that remains poorly studied in comparison to viruses and bacteria, but may pose considerable future risk of human transmissibility. Helminths are macroparasites, primarily known for chronic infections of the gastrointestinal tract, typically caused by tapeworms (cestodes), roundworms (nematodes), or flatworms (trematodes), although helminths can infect nearly all human tissues [17]. Helminths are also known to be vectors for other zoonoses, such as the fever-causing bacteria *Neorickettsia sennestu* transmitted by a trematode ingested via raw fish consumption [18]; although helminth vectoring remains understudied [19]. Human-helminth associations have ancient origins [reviewed in 20], but the relatively recent domestication of animals for food and companionship significantly increased the number of parasites shared between humans and (domesticated) animals [21]. The agricultural revolution and associated practices, such as storage of crops in granaries, likely created new links between humans and wildlife, providing additional opportunities for helminth species to infect human hosts [22]. To this day, zoonotic helminths continue to emerge within human populations, a process that may be further accelerated with the global trade of livestock, climate change and growth in the demand for animal protein for human consumption [23].

Helminths are distinct from other human parasites, such as viruses and bacteria, in that they commonly have complex life cycles that rely on one or more intermediate hosts [24, 25]. These intermediate hosts are necessary for the development of juvenile life stages (eggs and larvae) and transmission to the definitive host, where the animal matures, reproduces and produces propagules [26]. Intermediate hosts include a wide range of aquatic, terrestrial, wild and domesticated animals [26], yet it is unknown how intermediate host identities are linked to risk of helminthiasis in humans. In addition, transmission may occur directly (i.e., trophically, vertically) and/or indirectly (i.e., via environment or arthropod vector). From a public health perspective, most chronic infections are caused by soil-transmitted helminths [27], however, the transmission modes of most zoonotic helminths have not previously been summarized. Thus, identifying helminth biological and ecological traits that are linked to zoonosis can help to improve our understanding of the factors that drive zoonotic potential in helminths and to better manage risk of transmission to humans.

In addition to intrinsic biological and ecological traits such as identity of definitive and intermediate hosts, transmission to humans also may be influenced by socio-economic factors specific to regions where the parasites are found. Currently, most helminth infections in humans are found in low and middle income countries of the tropics [27, 28], where disease prevention and healthcare infrastructure vary greatly. Numerous parasitic worms such as hookworms (genera *Ancylostoma* and *Necator*) are considered neglected tropical diseases which could be eliminated with sufficient drug administration and effective interventions [28]. Further, given the generally high animal biodiversity of tropical regions, it also may be that there are more host species of potential zoonoses in this part of the world [29], although previous work indicates that temperate regions contain more zoonotic helminths than tropical regions [9]. Thus, we conjectured that geographic traits of helminths might be important factors for predicting the probability that a species might infect humans. Despite the high variation in medical, educational, and economic burden of human helminth infections worldwide [28], how the different epidemiological and geographic factors relate to helminth zoonotic potential has been unclear.

We investigated which traits of helminths are predictors of disease in humans. We compiled a global dataset from existing databases and the published literature on more than 700 mammal helminth parasite species to examine the frequency of biological (transmission, morphology), epidemiological, and geographical traits. We used boosted regression trees, an ensemble learning technique, to navigate the high dimensionality of these data. These and similar machine learning methods are rapidly developing approaches that can be applied to hetereogeneous covariates and are often robust to nonlinear interactions hidden in the data [30, 31]. Among over 70 variables, our machine learning approach identified key trait patterns predicting helminth zoonosis. Specifically, whether a helminth species is zoonotic was best predicted by three characteristics: (1) whether one of the hosts is a companion animal (i.e. dog, cat), (2) whether an intermediate host is a fish (member of Chordata phylum), and (3) the number of unique locations in which the helminth species has been detected. More generally, this study adds to the growing body of literature used to inform strategies for preventing helminth infection and mitigating risk of novel zoonoses.

## Methods

### Data compilation

We used the Global Mammal Parasite Database (GMPD) [32], which consists of over 700 species of helminths, representing three main phyla (Acanthocephala, Nematoda, and Platyhelminthes) of parasitic helminths that infect wild mammals. Most emerging zoonotic diseases originate from mammals [33] and therefore a mammal-focused analysis is well-suited to identifying zoonotic risk factors. For each helminth species, we searched primary literature for evidence of human infection originating from animal hosts to assign a binary response indicating whether or not the helminth species is zoonotic. We acquired morphological information of adults and eggs from Benesh, Lafferty [34] and Dallas, Gehman [35] databases, both of which gathered information from the literature. To fill in gaps, we followed Dallas, Gehman [35] and searched for missing morphological information from veterinary and parasitology references (e.g. Taylor, Coop [36]), taxonomy references [26, 37], and primary literature. We extracted minimum, mean, and maximum body length and width (in millimeters) of adult helminths from the descriptions of each parasite species. We also extracted minimum, mean, and maximum egg length and width (in millimeters). We compiled records of male and female body sizes when that information was available. We recorded site of infection in the definitive host body when it was provided.

We supplemented transmission information within the above references by extracting the following: common name(s) of definitive and intermediate hosts, whether the species has a free-living propagule stage (a binary variable), and if so, the stage of the free-living propagule as egg, larva, or both (as can occur in species that pass through more than one intermediate host), and the medium in which free-living stage(s) persist (soil, water, or both). We used the common names of intermediate hosts to note the class or phyla to which the intermediate animal host belongs, whether any of the host (definitive or intermediate) are domesticated animals (livestock and pets), or companion pet animals (predominantly cats and dogs). For each species we noted the transmission mode(s) to the definitive host as vertical (from parent to offspring), environmental (propagules acquired from the soil, water, or both), vector (via biting arthropod), or trophic (via consumption of intermediate host).

The GMPD provides geographical coordinates for each helminth species, which we augmented with host-helminth occurrence data from London Natural History Museum (LNHM) [38] available via R package helminthR [39]. Coordinates in the GMPD are from reported study site coordinates, or centroids of the reported study area [32]. Helminth occurrences in LNHM are georeferenced as centroids to the country or state (for the USA) level. In several instances coordinates were not provided by the databases, which we then georeferenced based on the location name using the *geocode* function [package ggmap; 40]. Some location names were obscure, such as the portion of a continent (e.g. southern South America) or body of water (e.g. southwest Atlantic), which we did not georeference. Next, based on the occurrence points of each species, we calculated the number of unique locations and latitudinal range (minimum and maximum), assigned a binary variable to indicate whether the species occurrences fall within the tropical latitudes (between 23° 27’ N and 23° 27’ S), and quantified the number of occurrences within tropical latitudes. We note that the number of unique locations reflects geographic distribution and sampling effort. From occurrence data we also calculated the number of countries, terrestrial ecoregions of the world (as defined by Olson, Dinerstein [41]), and terrestrial zoogeographic realms (as defined by Holt, Lessard [42]) from which each helminth species has been reported. Further, following Byers, Schmidt [43] we calculated range size for each helminth species as the total area of the ecoregions in which the species has been found. Finally, we calculated the mean gross domestic product (GDP) and human population size of the countries (provided by package rworldmap in R [44]) in which the species has been documented. Our final dataset consisted of 737 globally distributed helminth species (supplemental materials Fig. S1) and 73 trait variables describing helminth species that we included in our analyses. We classified the traits into one of four categories: transmission, epidemiological, morphological, or geographical traits (see Table 1). For full descriptions of each variable see supplementary materials.

**Table 1.** Top 15 most important variables used to predict helminth zoonoses status. Colors of the rows correspond to the four trait categories: geographical traits are in pink, transmission traits are in green, morphological traits are in blue and epidemiological traits are in orange. Color scheme also applies to Fig. 3 and 4.

| Variable | Description |
| --- | --- |
| **Transmission** |  |
| Pet host | Binary variable indicating whether the host (final or intermediate) is a companion animal (predominantly dog and cat) |
| Fish intermediate host | Binary variable indicating whether an intermediate host is a fish |
| **Geography** |  |
| Number of locations | Number of distinct locations (based on coordinates) a helminth species was observed in |
| Number of zoogeographic realms | Number of terrestrial zoogeographic realms (as defined in Holt et al 2013) a helminth species was located in |
| Number of tropical sites | Number of tropical sites the parasites was observed in |
| Number of ecoregions | Number of terrestrial ecoregions (as defined by World Wildlife Fund) |
| Number of countries | Number of countries the helminth parasite was observed in |
| **Morphology** |  |
| Male length (mean) | Mean male length in millimeters |
| Female length (max) | Maximum female length in millimeters |
| Egg width (max) | Maximum egg width in micrometers |
| Female length (min) | Minimum female length in millimeters |
| Male length (min) | Minimum male length in millimeters |
| Female width (max) | Maximum female width in millimeters |
| **Epidemiology** |  |
| Human population (mean) | Mean human population of the countries in which the helminth species is found |
| Gross domestic product (mean) | Mean gross domestic product (GDP) of the countries in which the helminth species occurs |

### Predictive model

We used boosted regression trees (BRT), a regression approach that permits missing data, variable interactions, collinearity, and non-linear relationships between the response and explanatory variables, which can be of mixed types [30, 45]. We fit a logistic-like predictive model with the zoonotic status of the helminths (0: not zoonotic, 1: zoonotic) as the response variable and the 73 traits as explanatory variables. Prior to analysis, we log transformed body size variables, which were right skewed. We randomly selected 80% of the data as the training set and reserved 20% for testing. Boosted regression trees were trained using the gbm package in R [46] with Bernoulli distributed error. We ran permutations of the model with different learning rates (1 × 10^−5^ to 1 × 10^−2^) and tree depths (1 to 3) using the training set to identify optimal learning parameters yielding the highest predictive performance (see supplementary materials Fig. S2). The learning conditions that were identified as yielding highest accuracy as assessed by the model AUC score (area under the receiver operating characteristic curve) included setting the maximum number of trees to 50,000, a learning rate of 0.001, and an interaction depth of 3. We used permutation procedures to compute relative importance scores for each predictor variable using Friedman’s algorithm [45]. We also build partial dependence plots, showing the marginal effect of each variable on the predicted outcome of the primary model [30, 45] (Fig. 1). Based on the results of the primary model, we ranked helminth species by their mean predicted probability of being transmissible to humans (Fig. 2).

**Figure 1.**
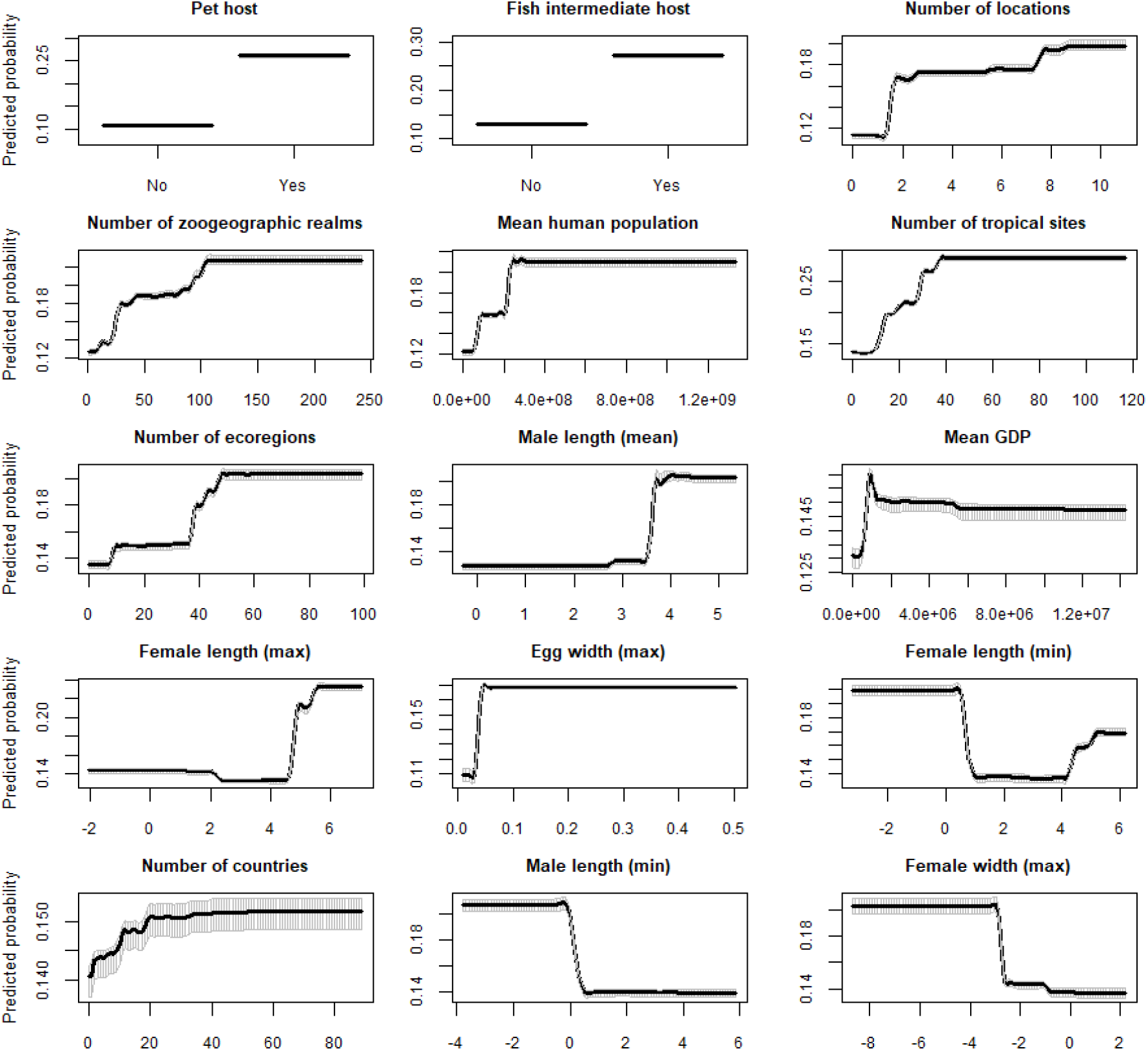
Partial dependence plots for top 15 most important variables. Plots are based on permutations of the primary boosted regression tree model that included 73 variables. Importance of the variables is ordered from left to right, then top to bottom. Black lines represent the median predicted probability, while shaded regions represent the corresponding 95% confidence interval across 100 permutations of the model.

**Figure 2.**
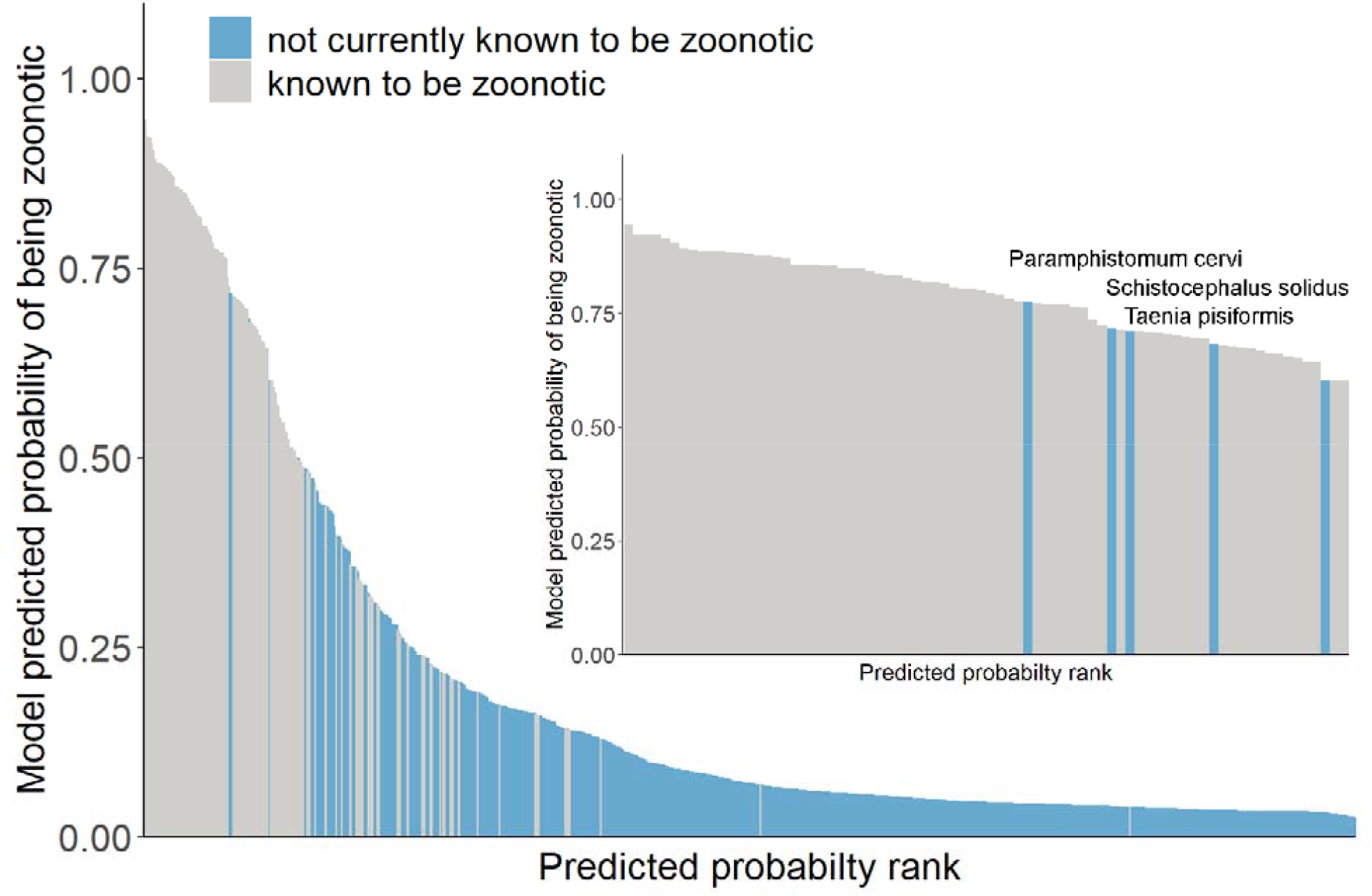
Predicted zoonotic helminth risk index. Average model-predicted probability of being zoonotic as ranked by the primary boosted regression tree model. Blue bars represent species not known to be transmissible to humans from wildlife and gray bars are species known to be transmissible to humans from wild hosts and are confirmed by the model to be zoonotic. Inset: zoonoosis risk of helminth species with model-predicted probabilities greater than 70%. Names of top 3 species not currently known to be zoonotic appear above the bars and include *Paramphistomum cervi, Schistocephalus solidus*, and *Taenia pisiformis* (in descending order).

Finally, we repeated the above analysis using only the top 15 most important variables predicted by the primary model trained on all 73 variables, and permuted the model 100 times. To further evaluate the relative importance of trait category, we ran additional submodels, also permuted 100 times, with one of the four trait categories (transmission, epidemiology, morphology, geography) excluded (Fig. 3 and 4). We used R programming for all analyses [47].

**Figure 3.**
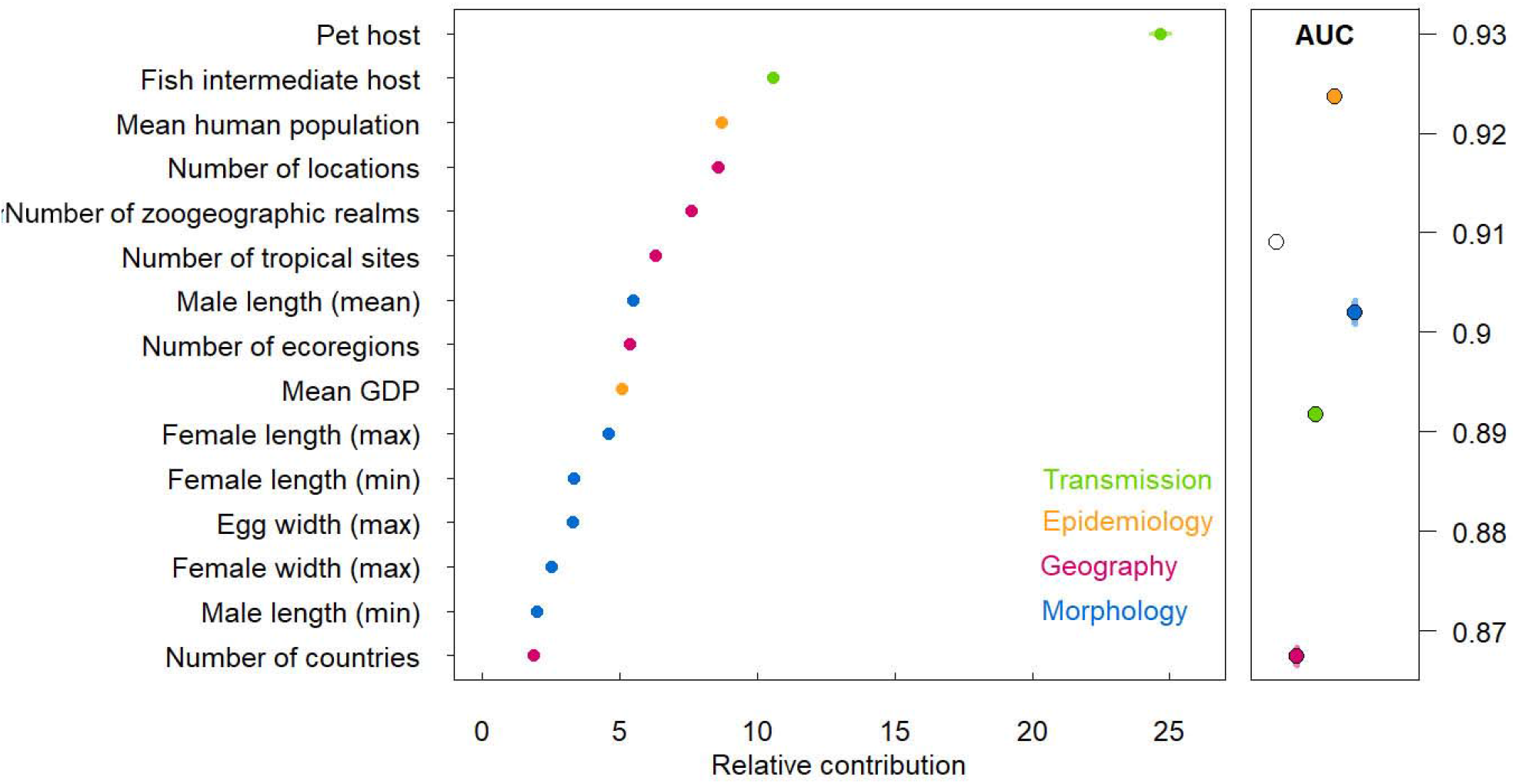
Variable importance values by permutation, averaged over 100 models trained on all four categories of traits (left panel), show relative importance of transmission traits (green), epidemiological traits (orange), geographical traits (maroon), and morphological traits (blue). Average model accuracy for each submodel trained on all four trait categories (white symbol), all trait categories except: morphological traits (blue), epidemiological traits (orange), transmission traits (green), or geographical traits (maroon). Error bars represent the standard deviation from 100 model permutations.

**Figure 4.**
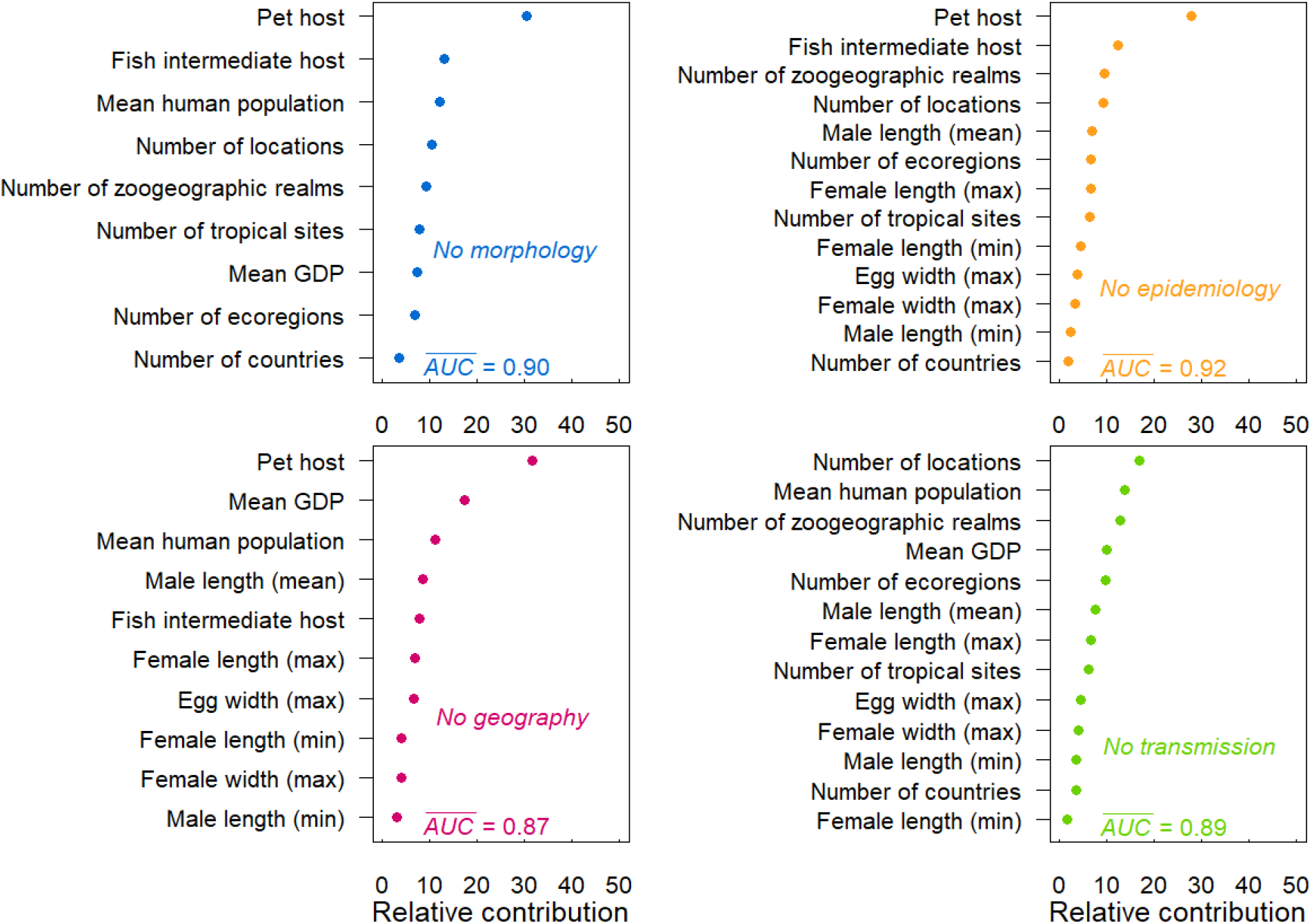
Variable importance values averaged over 100 model permutations trained on all categories of traits except: morphology (top left - blue), epidemiological traits (top right - orange), geography (bottom left - maroon), and transmission (bottom right- green).

## Results

We examined 737 globally distributed helminth species of which 137 are known to infect humans. Our boosted regression ensemble of models trained on 73 helminth traits distinguished zoonotic versus nonzoonotic species in the test dataset with 88% accuracy (AUC ± SE = 0.88 ± 0.01) and identified several predictors of zoonotic helminths (Fig. 1). The most important variable for accurately predicting zoonotic helminths was whether the helminth species is known to infect a companion animal, followed by whether fish serve as intermediate hosts, and the number of locations in which the helminth species has been documented. The fourth and fifth most important traits predicting zoonotic status in helminths related to the size of terrestrial zoogeographic regions observed for each helminth species (Fig. 1). Generally, the most important traits were related to geography and transmission, while epidemiological and morphological traits were least important (for the relative influence values of all 73 variables see supplementary materials Table S1).

While not currently known to cause human infection, BRT models identified 3 mammal-borne helminth species as likely to be zoonotic with >70% probability (Fig. 2) (in descending order): *Paramphistomum cervi, Schistocephalus solidus*, and *Taenia pisiformis*.

Additional ensembles of BRT models restricted to the top 15 most important variables (as identified by the primary models with 73 traits included, see Fig. 1) predicted the testing data with higher accuracy (AUC = 0.91) compared to the primary models trained on all 73 traits (AUC = 0.89). The restricted submodels trained on the 15 variables generally agreed on the ranking of the importance of variables with the primary models (Fig. 3). Submodels trained on data without one of the trait categories (i.e., leave-one-out) indicated that model trained on data without morphological traits performed slightly worse (AUC = 0.90) compared to submodels with all trait categories included (AUC = 0.91; Fig. 3), suggesting that including these features improved the predictive accuracy of our models. Models trained on data with epidemiological traits left-out performed best (AUC = 0.92; Fig. 3). Finally, models trained on data without geographical traits or transmission traits performed worse than models with other categories left out (AUC = 0.87, AUC = 0.89 respectively; Fig. 3). In submodels, companion animal host was the most important variable, except for the submodel that excluded transmission traits (Fig. 4). For AUC scores and the relative influence values of the variables in submodels see supplementary materials (Table S2).

## Discussion

Identifying pathogen traits associated with a propensity to spillover into humans is key for understanding and predicting emergence of novel human diseases originating from wildlife. We applied a machine learning algorithm to a large dataset of mammal helminths to identify characteristics distinguishing zoonotic and non-zoonotic species, and to predict which species currently classified as non-zoonotic have a high risk of ‘spilling over’ to humans in the future. Our results indicate that helminths that infect companion animals (dogs and cats) and utilize fish as intermediate hosts are more likely to cause human infection compared to other mammal-borne helminths. The third strongest predictor of the ability to cause human infection was the number of occurrences of helminth species, which indicates that widespread geographic distribution might provide important transmission exposure to human hosts; however, we note that this variable might also reflect sampling effort (see below). Overall, these results suggest that the zoonotic potential of helminth species is related to the identity of both definitive and intermediate hosts that come in direct and indirect contact with people, thereby providing abundant opportunities for parasite transmission. Further, our findings highlight the importance of transmission strategies in the ability of mammalian helminths to infect humans.

Particularly interesting is the predicted association between helminth zoonosis and companion animals (predominantly cats and dogs in this study). Domestic cats and dogs are hosts to numerous parasitic helminth species [36, 48] and represent an important link between humans and wildlife for zoonosis [49]. Indeed, the role of cats and dogs in helminthiasis have been well-documented for several parasites including the zoonotic tapeworm *Echinococcus multilocularis* (see Richards et al. in this issue) and roundworm *Toxocara cati* [49]. While many domesticated cats and dogs are “free-range” (i.e., not owned and cared for by humans), these animals are ubiquitous and tend to live near humans for provisioned food and shelter. Further, they hunt wild animals, consume animal parts (e.g. entrails) discarded by humans, and can overlap with wildlife habitat and territories [50], even in urban areas where numerous wild animals such as racoons, foxes, and coyotes thrive [51, 52]. The direct trophic interactions and indirect contacts dog and cats have with wildlife provide numerous opportunities for transmission of helminth parasites from wild to domestic animals, and eventually to humans. Additionally, the human-pet-wildlife interface has been around for centuries as it surfaced thousands of years ago with the domestication of cats 10,000 years ago and dogs 16,000 years ago [53, 54]. Therefore, there has been ample opportunity for host-jumping and host-switching events from wildlife to pets and humans, a process which is expected to accelerate with the increasing size of the human population, associated companion animals, and activities that impose close contact with wildlife.

Fish (freshwater or marine) as an intermediate host was identified as the third most important trait for predicting zoonosis. This finding is not surprising as fish are well-documented intermediate hosts to non-zoonotic parasitic worms that inflict humans [55]. One of the best-known examples of zoonotic parasites transmitted by fish is nematode *Anisakis simplex*, which have a complex life cycle with marine mammals as definitive host and high incidence among human populations that eat raw fish [56]. Fish-borne helminths are transmitted via consumption of raw, undercooked, or improperly preserved fish [57] and therefore fish represent an important direct trophic link between humans and wildlife. While *wild* fish are a source of parasitic helminths [55, 58], recent work indicates that farmed fish are also linked to zoonosis [59, 60]. Parasitic worm infections stemming from fish ingestion are increasing, likely due to the significant increase in demand for fish meat associated with changes in dietary habits and population growth [61]. Our finding elucidates fish as a key group of intermediate hosts linked to helminthiasis and the importance of monitoring fish intended for human consumption for parasitic worms to prevent and control zoonosis.

We also identified several geographical traits as important to predicting zoonotic helminths. Specifically, the number of unique locations around the world, the number of zoological realms in which helminths have been found, and the number of locations within the tropics were relatively important predictors. Overall, these findings suggest that mammalian parasitic helminths that are geographically widespread and able to persist in a range of habitat types are also more likely to be zoonotic than their more ecological specialized counterparts, possibly due to their ability to persist in different environmental conditions and exposure to humans in varying environments.

It is important to note that study effort (and attendant bias) is likely interwoven through several traits we included in this study. Particularly, the number of unique record locations might not only capture distribution but also number of samples and therefore sampling effort. Indeed, previous work shows that variation in sampling effort among parasitic species can predict the number of localities in which the species are documented [62]. Companion animal (pet host) trait might also reflect disproportionate study effort given the high access and relative ease of sampling. Furthermore, veterinary diagnostics (e.g. fecal floats, snap tests) more frequently performed on companion animals in high income countries might lead to higher discovery rate of helminth species in these places. We found that submodels which included or excluded the number of occurrences resulted in companion animal (pet host) remaining the most important predictor of zoonotic status among the helminths, lending some assurance of the strong statistical association between zoonotic status and pet host despite the influence of sampling effort in helminth data.

Our model predicted several helminth species that are currently not known to infect humans to have high probability (70% or higher) of causing zoonosis. The helminth species with highest probability of causing human infection was a flatworm, *Paramphistomum cervi*, followed by *Schistocephalus solidus*, and *Taenia pisiformis. Paramphistomum cervi* is environmentally transmitted and requires a snail intermediate host that is accidentally ingested by wild mammals and livestock ruminants (e.g. sheep and cattle), the definitive hosts [63]. Given that livestock can share species of gastrointestinal helminths with farmers [64], *Paramphistomum cervi* may be a likely candidate for spillover to humans. On the other hand, the flatworm *Schistocephalus solidus* infects a copepods, fish, and fish-eating water birds [65], all of which have the potential to provide trophic transmission to human host. *Taenia pisiformis* also appears likely to have the pathway to directly infect humans since it utilizes rabbit intermediate hosts and carnivores including cats and dogs as definitive hosts [66]. Indeed, consumption of wild rabbits by humans is popular in some European countries [e.g. Spain; 67] and might facilitate host-switching to humans for *Taenia pisiformis*. Identifying the three species of helminths and their traits serves as an initial step in focusing efforts on surveillance and empirical work investigating the zoonotic potential of these species.

In conclusion, we focused our study on parasitic helminth traits and used boosted regression trees to quantify how the different transmission, geographic, morphological and epidemiological factors relate to helminths’ zoonotic potential. Our work suggests that helminths found in cats and dogs are more likely to infect humans, and that consumption of fish by humans may pose a greater risk of spillover. While our study examined over 700 mammalian helminth species, many more parasitic worms are found in wildlife, and most are poorly described with little known about their life cycles [68]. Key life cycle details, such as intermediate host(s), are often assumed based on relation to better-studied species in the same genus. Large gaps in our understanding of life cycles and transmission dynamics exist for most parasitic worms, including those known to infect humans. Experimental infection work is largely lacking, while detailed studies of life cycles are no longer common [68] as molecular studies have eclipsed traditional experimental biology. Despite these knowledge gaps, the machine learning approach we took point to key insights about zoonotic helminths. In particular, our results highlight the importance of the interface between wildlife, companion animals, and humans in determining risk of parasitic worm infections, which continue to cause significant disease burden in developing countries [69], where semi-feral dogs and cats are generally not treated for parasites and will likely continue to serve as a source of novel helminthiases.

## Acknowledgements

We would like to thank J. Walker, C. Sanchez, and J. Vaz for helpful hints in compiling data and C. Cleveland for feedback on early version of the manuscript. We are also grateful to the UGA Library staff and Interlibrary Loan staff for the facilitating delivery of various articles for compilation of the trait data.

