## Supplementary materials for "Predictors of zoonotic potential in helminths"

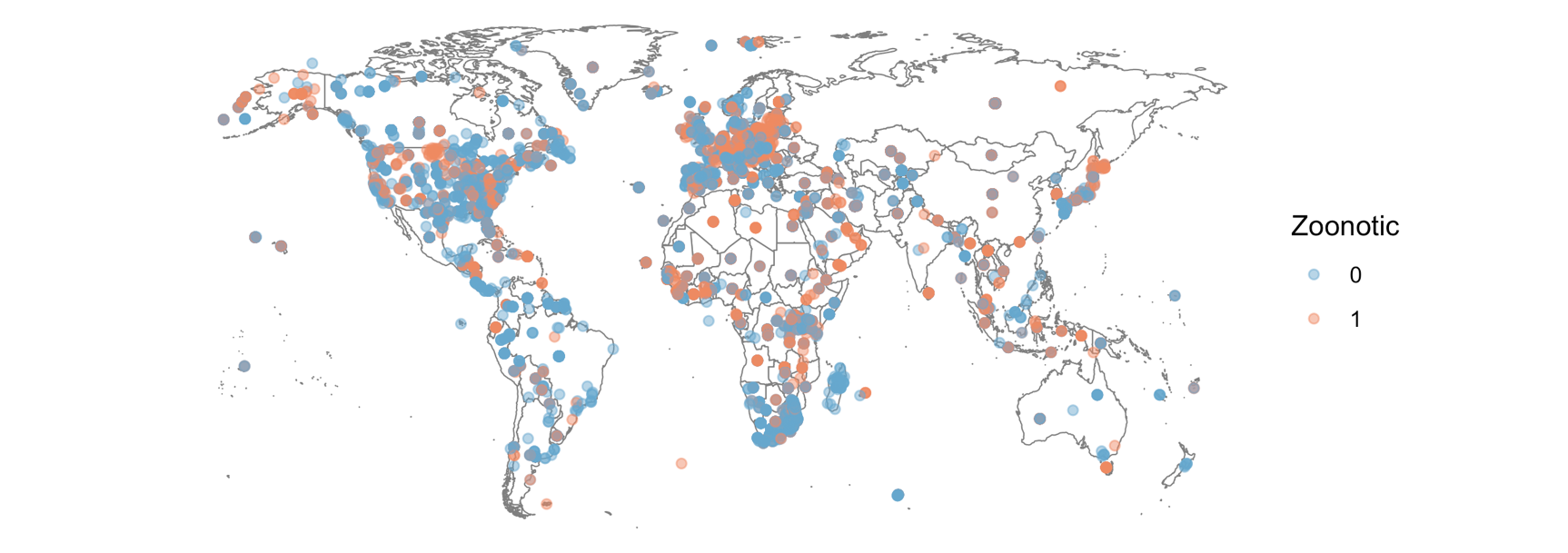

**Figure S1**. Distribution map of zoonotic (orange) and non-zoonotic (blue) helminth parasites occurrences. A total of 737 helminth species were utilized in this study.

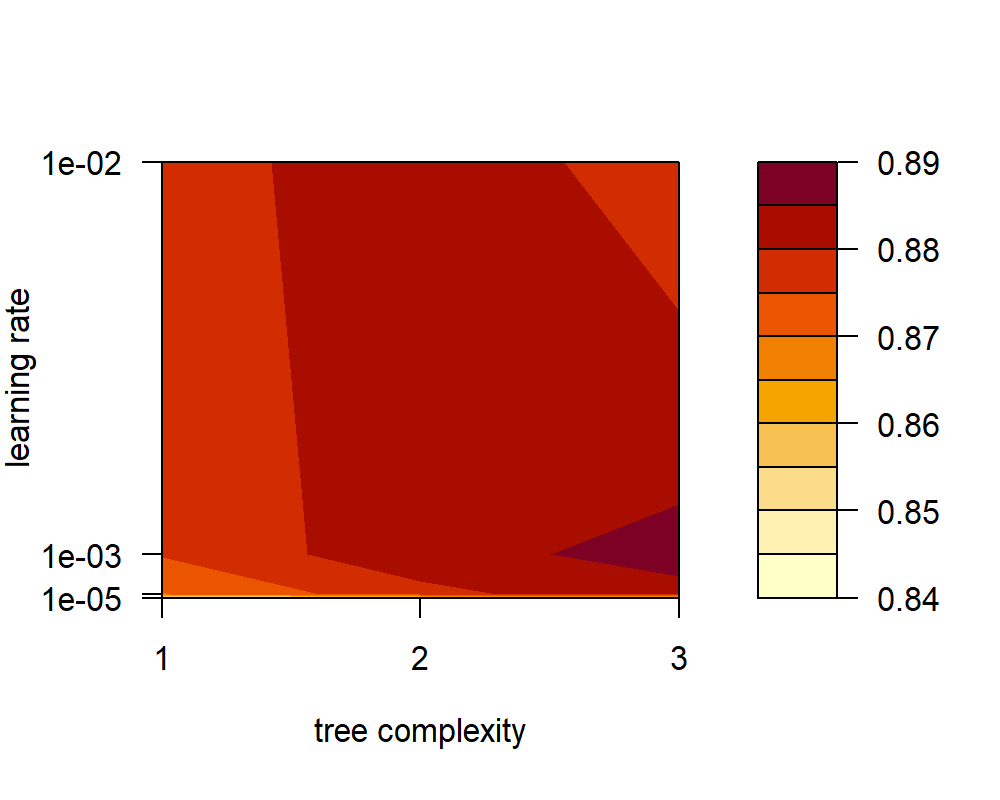

**Figure S2.** Contour plot of AUC scores resulting from boosted regression analyses with varying tree complexities and learning rates. Tree complexity of 3 and learning rate of 1e-03 results in highest AUC.

**Table S1. Relative importance of each of the 73 variables included in the full model ordered from highest to lowest importance.**

| **Variable** | **Influence** |
| --- | --- |
| Pet is a host | 24.64356 |
| Fish is an intermediate host | 9.492919 |
| Number of locations | 7.45715 |
| Number of zoological realms | 7.089547 |
| Mean human population | 6.681341 |
| Number of tropical sites | 5.551055 |
| Number of ecoregions | 5.304009 |
| Male length (mean) | 3.726184 |
| Mean GDP | 3.630745 |
| Female length (max) | 3.24017 |
| Egg width (max) | 2.705391 |
| Female length (min) | 2.640059 |
| Number of countries found in | 1.535795 |
| Male length (min) | 1.393169 |
| Male length (max) | 1.206644 |
| Free-living propagules persist in water | 1.116275 |
| Egg length (mean) | 1.083237 |
| Female width (max) | 1.052424 |
| Range size area | 0.961891 |
| Number of unique hosts identified | 0.936624 |
| Female length (mean) | 0.862995 |
| Total number of infection sites in definitive host body | 0.785656 |
| Trophic transmission | 0.740033 |
| Latitude (min) | 0.696618 |
| Latitude (max) | 0.666504 |
| Adult length (max) | 0.64263 |
| Adult width (min) | 0.461494 |
| Nonclose transmission | 0.36845 |
| Male width (min) | 0.355121 |
| Egg width (min) | 0.333846 |
| Infects the nervous system in definitive host | 0.307579 |
| Male width (max) | 0.257864 |
| Hosts are only wild (not domesticated) | 0.254421 |
| Adult width (max) | 0.242386 |
| Adult length (mean) | 0.23409 |
| Egg width (mean) | 0.209032 |
| Adult width (mean) | 0.144989 |
| Intermediate host is part of the lifecycle | 0.128008 |
| Free-living stage persist in soil | 0.121512 |
| Infects the skin in definitive host | 0.107101 |
| Female width (min) | 0.098517 |
| Infects the muscle in definitive host | 0.089761 |
| Egg length (max) | 0.073004 |
| Adult length (min) | 0.060348 |
| Livestock host | 0.055554 |
| Male width (mean) | 0.050572 |
| Female width (mean) | 0.040936 |
| Infects the digestive system in definitive host | 0.040029 |
| Environmental transmission | 0.036341 |
| Intermediate host in other class | 0.029633 |
| Amphibian intermediate host | 0.029381 |
| Crustacean intermediate host | 0.012704 |
| Eggs length (min) | 0.010373 |
| Maxillopod intermediate host | 0.00222 |
| Infects circulatory system in definitive host | 0.00211 |
| Close transmission | 0 |
| Infects the respiratory system in definitive host | 0 |
| Infects other systems in definitive host | 0 |
| Infects the reproductive system in definitive host | 0 |
| Reptilian is an intermediate host | 0 |
| Gastropod is an intermediate host | 0 |
| Aves is an intermediate host | 0 |
| Mammalia is an intermediate host | 0 |
| Insect is an intermediate host | 0 |
| Malacostraca is an intermediate host | 0 |
| Arachnid is an intermediate host | 0 |
| Vector transmission | 0 |
| Vertical transmission | 0 |
| Free-living stages do not pass via environment | 0 |
| Free-living stages occur in both soil and water | 0 |
| One of the hosts is wild animal | 0 |
| One of the hosts is a domestic animal | 0 |
| Present in the tropics | 0 |

**Table S2. Relative importance of top 15 variables included in submodels along with AUC scores.**

|  | **Relative importance** | | | | |
| --- | --- | --- | --- | --- | --- |
| **Variable** | **All included** | **No morphology** | **No epidemiology** | **No geography** | **No transmission** |
| Pet host | 24.63 | 30.55 | 27.11 | 30.40 | not included |
| Fish intermediate host | 10.55 | 13.01 | 12.07 | 7.63 | not included |
| Mean human population | 8.72 | 12.10 | not included | 11.27 | 13.59 |
| Number of locations | 8.49 | 10.28 | 9.37 | not included | 16.88 |
| Number of zoogeographic realms | 7.52 | 9.21 | 9.31 | not included | 12.90 |
| Number of tropical sites | 6.29 | 7.58 | 6.37 | not included | 6.05 |
| Male length (mean) | 5.48 | not included | 6.69 | 8.29 | 7.63 |
| Number of ecoregions | 5.33 | 6.76 | 6.55 | not included | 9.63 |
| Mean GDP | 5.12 | 6.95 | not included | 17.54 | 9.84 |
| Female length (max) | 4.69 | not included | 6.64 | 6.92 | 6.50 |
| Female length (min) | 3.37 | not included | 4.50 | 3.93 | 1.69 |
| Egg width (max) | 3.32 | not included | 3.75 | 6.52 | 4.44 |
| Female width (max) | 2.54 | not included | 3.38 | 4.29 | 3.94 |
| Number of countries | 1.99 | 3.56 | 1.92 | not included | 3.36 |
| Male length (min) | 1.97 | not included | 2.36 | 3.21 | 3.55 |
|  | **AUC** | | | | |
|  | 0.91 | 0.90 | 0.92 | 0.87 | 0.89 |

**Table S3:** Variable description and definitions for data used in the study. Data and accompanying code is available on figshare.

| **Variable** | **Definition** |
| --- | --- |
| Parasite | Helminth species name as found in GMPD 2.0 |
| ZooScore | Zoonosis score: 0 = in humans and wildlife but not acquired through vetebrate reservoir; 1 = zoonotic and can be acquired through vetebrate reservoir; 2 = zoonotic but no transmissible to other humans;  3 = zoonotic and transmissible to other humans |
| ConfidScore_ZooScore | Confidence for ZooScore: 1= most confident, 2 = somewhat confident, 3 = least confident |
| close | Parasite transmissible via close non-sexual contact such as  grooming, biting, scratching, aerosols. |
| nonclose | Parasite transmissible via non-close contact such as by fomites or ingestion of food or water contaminated with feces or urine. |
| intermediate | Intermediate Parasite transmitted by intermediate hosts such as snails or crustaceans (complex life cycles) |
| ParType | Parasite type. All rows are “Helminth” |
| ParPhylum | Parasite Phylum |
| ParClass | Parasite Class |
| ParOrder | Parasite Order |
| ParFamily | Parasite Family |
| intermediate.hosts | Common or vernacular name of the intermediate hosts |
| paratenic.host | Common name of the paratenic hosts. 'Paratenic' indicates a host in which parasite can cause infection or be present but no development of the stage occurs. |
| typical.definitive.hosts | Typical host(s) for the parasite |
| sites.of.infection | Where the helminth occurs in the host |
| migration.to.tissue | What tissues the helminth travel through |
| transmission.mode.to.final.host | Transmission mode to the definitive host |
| TM.confidence | Confidence score of the transmission mode. 3 = transmission experimentally well studied; 2 = some parts of transmission are studied known but some parts are assumed based on other well known related species; 1= little known, mostly assumed based on related species; 0 = no information found |
| infection.type.in.humans..Chronic.Acute. | Whether the parasite causes chronic or acute human infection |
| medium.of.infective.stages.between.hosts..soil..water. | Medium in which free living stages are found (soil, water, or both) |
| exter.life.stage | External infective stage (egg, larva, or both) |
| dom.host.name | Common name of domestic animal host (intermediate or definitive) |
| daily.egg.larva.production | Number of eggs or larvae produced per day |
| longevity.of.free.living.stage..days. | Number of days free living stages survive in the medium |
| prepatent.period..d. | Number of days between infection and the recovery of an infective stages from the blood or feces. |
| merged.syn | Yes or no whether the species consists of multiple merged species formerly considered separate species |
| host_count_B | Number of hosts based on Benesh et al. 2016 |
| FemaleMeanLength | Mean female length in millimeters |
| FemaleMaxLength | Maximum female length in millimeters |
| FemaleMeanWidth | Mean female length in millimeters |
| FemaleMaxWidth | Maximum female width in millimeters |
| FemaleMinLength | Minimum female length in millimeters |
| FemaleMinWidth | Minimum female width in millimeters |
| MaleMeanLength | Mean male length in millimeters |
| MaleMaxLength | Maximum male length in millimeters |
| MaleMeanWidth | Mean male length in millimeters |
| MaleMaxWidth | Maximum male width in millimeters |
| MaleMinLength | Minimum male length in millimeters |
| MaleMinWidth | Minimum male width in millimeters |
| AdultMeanLength | Mean adult length in millimeters |
| AdultMaxLength | Maximum adult length in millimeters |
| AdultMeanWidth | Mean adult length in millimeters |
| AdultMaxWidth | Maximum adult width in millimeters |
| AdultMinLength | Minimum adult length in millimeters |
| AdultMinWidth | Minimum adult width in millimeters |
| EggsMeanLength | Mean egg length in millimeters |
| EggsMaxLength | Maximum egg length in millimeters |
| EggsMeanWidth | Mean egg length in millimeters |
| EggsMaxWidth | Maximum egg width in millimeters |
| EggsMinLength | Minimum egg length in millimeters |
| EggsMinWidth | Minimum egg width in millimeters |
| minLat | Minimum latitude of occurrence |
| maxLat | Maximum latitude of occurrence |
| meanHumPop | Mean human population of the countries in which the parasite is found |
| meanGDP | Mean GDP of the countries in which the parasite is found |
| NUMtropicSites | Number of tropical sites the parasites is observed in |
| numCountries2 | Number of countries the helminth parasite was observed in |
| numCMECRealms | Number of terrestrial zoogeographic realms helminth parasite was located in as defined in Holt et al 2013 (DOI: 10.1126/science.1228282) |
| numEcoRegions | Number of terrestrial ecoregions as defined by Olson et al. (2001) (layer, 2011; Source data available here: http://maps.tnc.org/gis_data.html). There are 827 ecoregions. |
| loc_count | Number of distinct location (based on lat and long) the helminth parasite was observed in |
| TNC_SHAPE_AreaSUM | Parasite range size area equal to total area of the ecoregions in which the species has been found |
| digestive | Binary variable indicating whether site of infection in the host body consisted of digestive tract |
| locomot | Binary variable indicating whether site of infection in the host body consisted of muscles |
| respir | Binary variable indicating whether site of infection in the host body consisted of respiratory tract |
| inf.other | Binary variable indicating whether infection site is other than digestive, locomotor, respiratory, nervous, skin, or reproductive system |
| nervous | Binary variable indicating whether site of infection in the host body consisted of the nervous system |
| skin | Binary variable indicating whether site of infection in the host body consisted of epidermal tissue |
| circul | Binary variable indicating whether site of infection in the host body consisted of circulatory system |
| reprod | Binary variable indicating whether site of infection in the host body consisted of reproductive organs and tissues |
| total.inf.sites | Total number of sites of infection in the host body |
| IH_Amphibia | Binary variable indicating whether an intermediate host is amphibian |
| IH_Crustacea | Binary variable indicating whether an intermediate host is crustacean |
| IH_Reptilia | Binary variable indicating whether an intermediate host is reptilian |
| IH_Gastropoda | Binary variable indicating whether an intermediate host is gastropod |
| IH_Aves | Binary variable indicating whether an intermediate host is a bird |
| IH_Mammalia | Binary variable indicating whether an intermediate host is mammalian |
| IH_Insecta | Binary variable indicating whether an intermediate host is an insect |
| IH_Malacostraca | Binary variable indicating whether an intermediate host is member of class Malacostraca |
| IH_Chordata | Binary variable indicating whether an intermediate host is member of class Chordata (i.e., fish) |
| IH_Maxillopoda | Binary variable indicating whether an intermediate host is member of the class Maxillopoda |
| IH_Arachnida | Binary variable indicating whether an intermediate host is a member of the class Arachnida |
| IH_other | Binary variable indicating whether an intermediate host is of other class |
| Environmental | Binary variable indicating whether a transmission mode is environmental |
| Trophic | Binary variable indicating whether a transmission mode is trophic |
| Vector | Binary variable indicating whether a transmission mode is vectored by biting arthropod |
| Vertical | Binary variable indicating whether a transmission mode is vertical |
| wild_only | Binary variable indicating whether hosts are wild species only |
| soil_med | Binary variable indicating whether propagule transmission medium is soil |
| water_med | Binary variable indicating whether propagule transmission medium is water |
| no_med | Binary variable indicating whether propagule is transmitted via medium |
| both_s_w | Binary variable indicating whether transmission medium is both soil and water |
| LivestockH | Binary variable indicating whether host is a livestock animal |
| PetH | Binary variable indicating whether host is a companion animal |
| WildH | Binary variable indicating whether host is a wild animal |
| DomesticH | Binary variable indicating whether host is a domesticated animal |
| Tropical | Binary variable indicating whether the species occurs in the tropics |
| Zoonotic | Binary variable indicating whether species is zoonotic |
| Notes | Additional notes on the species |
| References | List of additional references for the species |
| DataSources | List of data sources used to compile traits |
